# Population genomics of nicotinic acetylcholine receptors in *Anopheles funestus* reveals rapid evolution of the α9 and β2 subunits within a constrained gene family

**DOI:** 10.64898/2026.05.15.725454

**Authors:** Desiree E. Rios, Caroline Fouet, Colince Kamdem

## Abstract

The deployment of clothianidin-based insecticide formulations in malaria vector control has highlighted the capacity of *Anopheles funestus* to displace more susceptible mosquito species in treated areas and to rapidly evolve resistance under selection pressure. Metabolic detoxification, together with structural and genetic changes in nicotinic acetylcholine receptors (nAChRs), the primary molecular targets of neonicotinoids, can reduce insecticide efficacy. Here, we characterized amino acid substitutions across all 11 nAChR subunits in *An. funestus* to assess standing variation that may facilitate adaptive responses to chemical exposure. Using whole-genome sequencing data from 656 mosquitoes sampled in 13 African countries, we found marked contrasts in the distribution of nonsynonymous variants among nAChR subunits. Most subunits are strongly constrained and carry no missense variants, whereas two loci (α3 and α7) display three geographically widespread amino acid substitutions across the continent. In contrast, α9 and β2 accumulate dozens of nonsynonymous mutations occurring at intermediate to high frequencies, including within domains involved in orthosteric ligand binding and channel gating. Genetic differentiation at nAChR loci among populations from different countries is low to moderate, although several nonsynonymous mutations display high *F*_ST_ values consistent with geographic structuring. These results highlight relaxed constraint on two subunits that may provide opportunities for evolutionary diversification within a conserved family of multimeric receptor assemblies. Such diversification has not been observed in vector species displaced by *An. funestus* in indoor residual spraying areas, and the potential implications for reduced sensitivity to neonicotinoids are discussed.

## Introduction

Clothianidin-based insecticide formulations have been introduced into malaria vector control programs to mitigate widespread resistance to pyrethroids (1–3). However, their deployment has also revealed rapid shifts in vector species composition, with less susceptible species replacing more sensitive populations in treated areas. In Uganda, field studies reported the replacement of *Anopheles gambiae* by *An. funestus* within two years of indoor residual spraying with clothianidin-based formulations, concomitant with the rapid emergence of resistance in *An. funestus* (4,5). When clothianidin was subsequently replaced with the organophosphate pirimiphos-methyl, *An. gambiae* re-emerged, and follow-up assays confirmed that the observed species shift was associated with the lower sensitivity of *An. funestus* to clothianidin. These observations are consistent with laboratory studies indicating substantial differences in baseline susceptibility to neonicotinoids among *Anopheles* species, including between closely related taxa (6–8).

Cross-resistance with other insecticide classes, such as pyrethroids, may contribute to the capacity of *An. funestus* to develop resistance to neonicotinoids (9). However, the distinct modes of action of neonicotinoids and pyrethroids reduce the likelihood of direct cross-resistance between these insecticide classes, as confirmed by mortality assays in adult *Anopheles* (1,10,11). Cytochrome P450-mediated detoxification represents one of the principal mechanisms by which insect pests evolve reduced sensitivity to neonicotinoids (8,12–17). Contemporary populations of *An. funestus* are characterized by overexpression of multiple cytochrome P450 enzymes (CYP), some of which may contribute to neonicotinoid metabolism (18–20). Recent studies suggest that some CYPs, including the duplicated *CYP6P9a* and *CYP6P9b*, may reduce *An. funestus* sensitivity to clothianidin, although direct in vitro evidence of their metabolic activity against neonicotinoids is still lacking (21). In addition, synergist bioassays indicated that mortality is not fully restored in the presence of the CYP inhibitor piperonyl butoxide, which suggests that CYP-mediated metabolism alone may not fully account for reduced clothianidin susceptibility in mosquitoes (7,22,23). These findings imply that additional mechanisms, such as alterations in insecticide target sites, may act in combination with enhanced detoxification capacity to reduce *An. funestus* susceptibility to clothianidin.

In insects, nicotinic acetylcholine receptors (nAChRs) are the primary molecular targets of neonicotinoids, as well as of related insecticide classes such as spinosyns and butenolides (24–26). As members of the Cys-loop ligand-gated ion channel superfamily, nAChRs are pentameric transmembrane proteins whose activation by the neurotransmitter acetylcholine mediates fast cholinergic neurotransmission in the central nervous system (27). nAChRs are essential for synaptic transmission and play a key role in regulating multiple behavioral and physiological processes in insects, including sleep, circadian rhythms, escape behavior, and immunity (28–33). Insect nAChRs are encoded by a relatively small gene family comprising 10-12 α and non-α subunits (34). Sequence divergence among subunits, together with variation in subunit composition, generates substantial functional diversity in the pentameric receptors (35–39).

Neonicotinoids generally are agonists that mimic acetylcholine at the orthosteric binding site of nicotinic acetylcholine receptors (nAChRs) (24,40). Binding of neonicotinoids results in sustained receptor activation, leading to neuronal overstimulation, paralysis, and death in insects. In each pentameric receptor, the orthosteric binding site is located at the extracellular interface between adjacent subunits and is defined by a set of conserved binding loops (41–43). Ligand binding involves electrostatic interactions and hydrogen bonding between canonical amino acid residues located in loops A-C or G on the principal face of an α subunit and loops D-G on the complementary face of a neighboring non-α subunit, or another α subunit in homomeric assemblies (24,40,44).

Extensive mutagenesis and structural studies have identified conserved residues within the orthosteric binding loops that determine neonicotinoid affinity (45–52). In contrast to mammals and ticks, whose nAChRs exhibit lower affinity for neonicotinoids, insect receptors contain basic residues within loops that enable stable binding of active ingredients (46,53,54). Mutations at these positions, particularly within the β1 subunit, can substantially reduce binding affinity and have been linked to resistance development in several agricultural pests (55–59). In addition to target-site mutations, the differential expression of nAChR subtypes with intrinsically lower affinity for neonicotinoids also contributes to reduced insecticide sensitivity in insects (60–62).

Beyond changes in ligand-binding affinity, Cys-loop receptors contain multiple agonist-binding sites, noncompetitive antagonist sites, and gating elements that interact at a distance through conformational changes in the receptor’s quaternary structure (42). These sites provide interfaces through which xenobiotic compounds can modulate allosteric changes, ion selectivity, post-translational modifications, and interactions with membrane lipids (63–67). Consequently, systematic investigation of genetic variation within the nAChR gene family is essential for identifying residues involved in receptor diversification and potentially in neonicotinoid sensitivity. With the growing availability of high-quality whole-genome sequencing data for vector species, including *An. funestus*, such analyses are now within reach (68–71).

The objective of this study was to use whole-genome sequencing data to characterize amino acid substitutions in nAChRs in wild populations of the major malaria vector *An. funestus*. Although the nAChrR gene family is strongly constrained, relaxed purifying selection enables the accumulation of intermediate- to high-frequency nonsynonymous variants in the α9 and β2 subunits, including within canonical ligand-binding loops and transmembrane domains. In comparison, the major malaria vectors *An. gambiae* and *An. coluzzii*, which are more susceptible to neonicotinoids than *An. funestus*, harbor only rare nonsynonymous variants across the nAChR gene family (72). Geographically widespread standing variation on divergent subunits may provide the evolutionary resources facilitating *An. funestus* responses to neonicotinoid exposure.

## 2. Materials and Methods

### 2.1 Mosquito populations and whole-genome dataset

We analyzed genetic variation within the nAChR gene family using whole-genome sequences from 656 wild-caught *An. funestus* mosquitoes sampled across 13 sub-Saharan African countries. The dataset has been described in detail previously (69). Adult mosquitoes were collected between 2014 and 2018, largely before the deployment of clothianidin-based indoor residual spraying in Africa (4,5). The 656 individuals included in this study were sequenced at a minimum median coverage of 10×. Variant call format (VCF) files containing genotype information for individual mosquitoes were used for downstream analyses (69).

### 2.2 Identification of nAChR subunits in the *An. funestus* genome

We identified genes encoding nAChR subunits (α and β) in the *An. funestus* genome assembly (idAnoFuneDA-416_04 updated to AfunGA1) using orthologous sequences from the *An. gambiae* PEST reference genome (35,72,73). Each *An. gambiae* subunit and the corresponding ortholog in the *An. funestus* genome assembly (AfunGA1) were both retrieved from VectorBase (74). The genomic positions of the 11 subunits were compared between species to assess conservation of chromosomal location and synteny. Protein sequences of orthologous subunits were aligned using MAFFT (75) to estimate sequence similarity between species. A multiple sequence alignment of all *An. funestus* nAChR subunits was then performed to characterize conserved structural features within species. This alignment enabled the mapping of conserved Cys-loop receptor features, including ligand-binding loops (A–G) and transmembrane domains (TM1–TM4) as well as the discrimination of α and β subunits based on the presence of the characteristic adjacent cysteine residues in loop C. Domain boundaries were identified based on conserved sequence motifs described previously (35).

### 2.3 Variant detection and annotation in *An. funestus* nAChRs

To detect single-nucleotide polymorphisms (SNPs) in *An. funestus* nAChRs, we used the genomic coordinates of each subunit together with bcftools v1.21 (76) to extract variants from VCF files generated for the 656 sequenced individuals. For each gene, a merged VCF file containing all intronic and exonic SNPs was generated and used as input for variant annotation. Variant annotation was conducted with SnpEff v4.3 (77), which classified mutations according to their predicted functional impact within each nAChR gene. The *An. funestus* database used for SnpEff annotation was generated from the reference genome (AfunGA1) and its corresponding GFF file. Annotated VCF files were filtered to retain variants classified as either “high impact,” corresponding to predicted loss-of-function mutations, “moderate impact,” indicating missense variants, and “low impact” encompassing all synonymous mutations. Variants with more than 20% missing genotypes across samples were excluded. Annotation fields were extracted from filtered VCF files using SnpSift (78). Only mutations identified on a single transcript and classified as high-, moderate- or low impact by SnpEff were retained for downstream analysis.

### 2.4 Allele frequencies, genetic differentiation, and structural localization of amino acid substitutions

We calculated frequencies for each variant within the 13 sampled countries and across the continent using PLINK v2.0 (79,80). Variants were then assigned to frequency classes to create allele frequency spectra. Variants were classified as: low (0.05–0.25), intermediate (0.25–0.50), and high (0.50–1.00). Genetic differentiation among populations from different countries was quantified for each nonsynonymous variant with allele frequency > 0.05 using Hudson’s *F*_ST_ estimator implemented in PLINK v2.0. In each nAChR subunit, pairwise *F*_ST_ values were calculated for all country combinations, and median *F*_ST_ was used to measure overall differentiation. Genetic differentiation at each nonsynonymous mutation was classified as low (median *F*_ST_ < 0.05), moderate (0.05-0.15), high (0.15-0.25), or very high (> 0.25). To identify highly differentiated variants within each subunit, we used the 95^th^ percentile of the distribution of pairwise *F*_ST_ values as a threshold.

To examine the distribution of amino acid substitutions within nAChR pentameric assemblies, we generated structural models for specific receptor subtypes. We restricted this analysis to subunits exhibiting more than four nonsynonymous mutations at allele frequencies > 0.05 (α9 and β2). We used α1 as a complementary subunit to create a heteropentamer containing β2: (α1)₃(β2)₂. Likewise, to generate a heteropentamer containing α9, we used β1 as a complementary subunit to construct the (α9)₃(β1)₂ receptor subtype. AlphaFold2 models for *An. funestus* α1 and β1 were obtained from the AlphaFold database, whereas structural models for α9 and β2 were created in ColabFold using the protein sequence as input (81). To further characterize the orthosteric ligand-binding domain, we used the human α7 nAChR structure with bound agonist Epibatidine (EPB) to delineate the ligand-binding pocket. The human α7 nAChR with bound EPB was retrieved from the Protein Data Bank (PDB ID: 7EKP) and used as a template to model the pentameric structure of *An. funestus* subtypes.

Structural alignment between the human α7 template and *An. funestus* subunits was performed in PyMOL (82), yielding alignments with root-mean-square deviation (RMSD) < 2 Å. α and β subunits aligned to the human α7 template with the bound EPB ligand were combined to generate the desired heteropentameric models: (α1)₃(β2)₂ and (α9)₃(β1)₂. Naturally occurring amino acid substitutions identified in *An. funestus* α9 and β2 were mapped onto these structural models to visualize their spatial distribution relative to ligand-binding pockets and transmembrane domains.

## 3. Results

### 3.1. High sequence similarity between *An. funestus* and *An. gambiae* nAChRs

All 11 nAChR subunits identified in *An. gambiae* were present in *An. funestus* (Table 1). Subunits located on the X chromosome in *An. gambiae* (α3, α4, α7, and β1) were also found on the X chromosome in *An. funestus*. Four subunits situated on chromosome 2R in *An. gambiae* (α1, α2, α6, and α8) were found on the corresponding 2RL arm in *An. funestus*. In contrast, three subunits located on chromosome 3R in *An. gambiae* (α5, α9, and β2) were mapped to chromosome 2RL in *An. funestus* (Table 1).

**Table 1.** Genomic location of nicotinic acetylcholine receptor (nAChR) subunits in *Anopheles gambiae* and *An. funestus*.

| Subunits | *An. gambiae* |  | *An. funestus* |  |  |  |  |  |
| --- | --- | --- | --- | --- | --- | --- | --- | --- |
|  | Accession number | Chromosome | Accession number | Chromosome | Transcript | Start | Stop | Strand |
| $\alpha 1$ | AGAP002974 | 2R | AFUN2006507 | 2RL | R10788 | 9,997,743 | 10,051,521 | + |
| $\alpha 2$ | AGAP002972 | 2R | AFUN2007949 | 2RL | R13654 | 9,917,926 | 9,954,863 | - |
| $\alpha 3$ | AGAP000329 | X | AFUN2006724 | X | R11231 | 4,862,865 | 4,873,400 | + |
| $\alpha 4$ | AGAP000138 | X | AFUN2003601 | X | R4927 | 14,811,911 | 14,837,423 | - |
| $\alpha 5$ | AGAP008588 | 3R | AFUN2002573 | 2RL | R2897 | 66,353,520 | 66,426,071 | - |
| $\alpha 6$ | AGAP002152 | 2R | AFUN2010718 | 2RL | R19105 | 10,462,426 | 10,478,752 | + |
| $\alpha 7$ | AGAP000962 | X | AFUN2006065 | X | R9936 | 14,468,113 | 14,549,127 | - |
| $\alpha 8$ | AGAP002971 | 2R | AFUN2014450 | 2RL | R26683 | 9,876,310 | 9,931,054 | + |
| $\alpha 9$ | AGAP009493 | 3R | AFUN2006311 | 2RL | R10413 | 97,489,370 | 97,493,281 | + |
| $\beta 1$ | AGAP000966 | X | AFUN2001135 | X | R28201 | 14,589,007 | 14,596,533 | + |
| $\beta 2$ | AGAP010057 | 3R | AFUN2008562 | 2RL | R14912 | 61,381,665 | 61,383,773 | + |

Pairwise protein alignments between *An. gambiae* and *An. funestus* orthologs revealed high sequence similarity for all subunits (95.75% on average). Amino acid identity ranged from 72.9% (β2) to 99.8% (α1) (Table 2). Multiple sequence alignment of all 11 *An. funestus* subunits confirmed the presence of conserved structural elements characteristic of Cys-loop pentameric ligand-gated ion channels (Fig. 1). These included seven ligand-binding loops (A–G) containing canonical residues involved in acetylcholine and neonicotinoid binding, four conserved transmembrane domains (TM1–TM4), and the adjacent cysteine residues in loop C that distinguish α from β subunits. The region with the greater variability among *An. funestus* subunits corresponded to the intracellular loop between TM3 and TM4.

**Figure 1.**
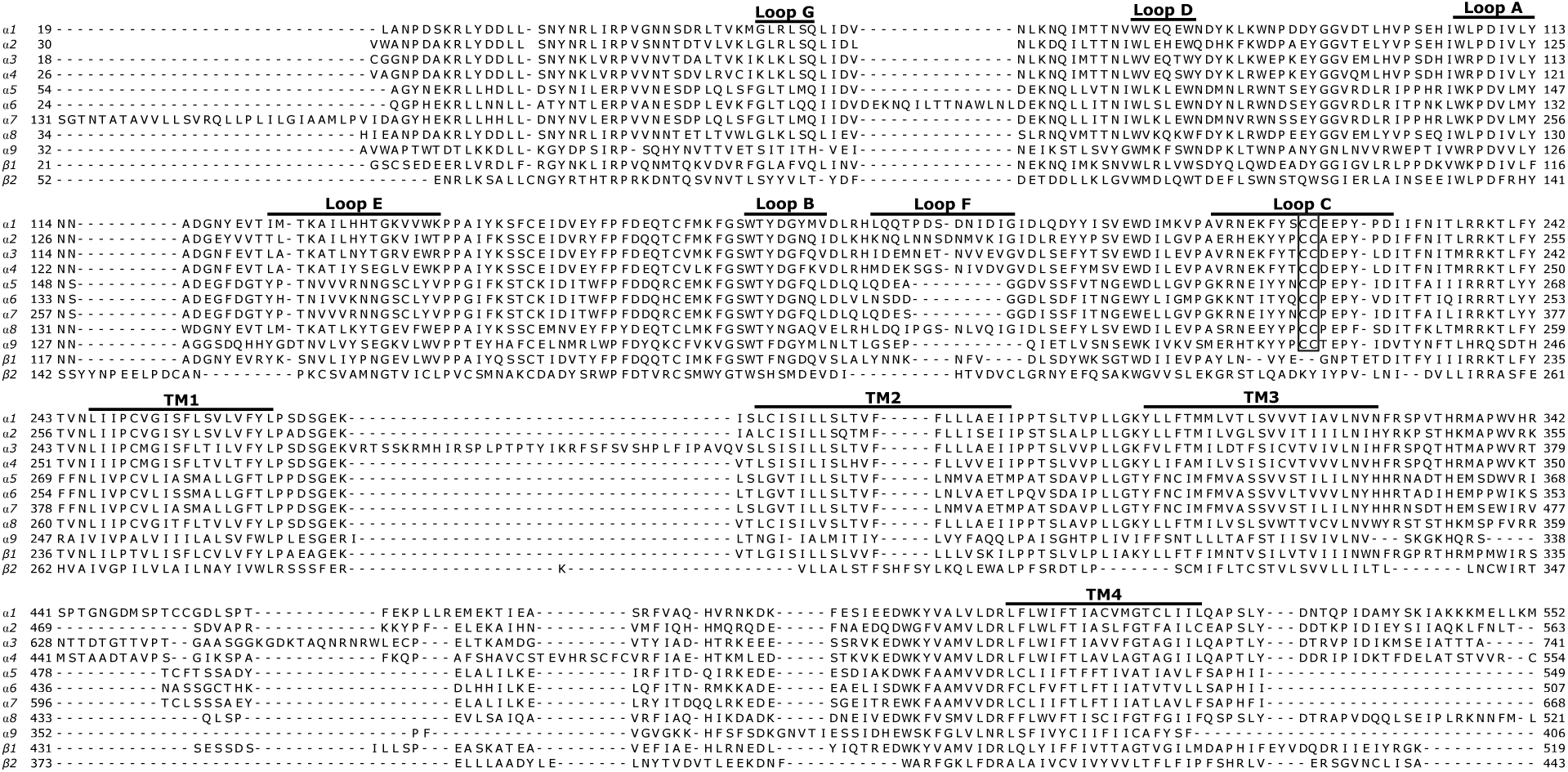
Multiple sequence alignment of nicotinic acetylcholine receptor (nAChR) subunits in *An. funestus*. Conserved features of Cys-loop ligand-gated ion channels are highlighted, including the extracellular ligand-binding loops (A–G) and the four transmembrane domains (TM1–TM4). The adjacent cysteine residues in loop C that define α subunits are indicated by a box.

**Table 2.** Sequence comparison of nAChR subunits between *An. gambiae* and *An. funestus*.

| Subunits | $\alpha 1$ | $\alpha 2$ | $\alpha 3$ | $\alpha 4$ | $\alpha 5$ | $\alpha 6$ | $\alpha 7$ | $\alpha 8$ | $\alpha 9$ | $\beta 1$ | $\beta 2$ |
| --- | --- | --- | --- | --- | --- | --- | --- | --- | --- | --- | --- |
| Length (number of amino acids) | 557 | 569 | 754 | 557 | 552 | 509 | 676 | 531 | 406 | 519 | 446 |
| Identity (%) | 99.8 | 98.9 | 91.2 | 97.1 | 99.6 | 99.2 | 86.5 | 83.8 | 88.9 | 99.4 | 72.9 |
| Similarity (%) | 100 | 99.5 | 91.5 | 98.9 | 99.8 | 99.6 | 89.2 | 83.8 | 95.1 | 99.6 | 84.3 |

### 3.2. Extensive amino acid substitutions across the α9 and β2 subunits

Analysis of 656 mosquitoes sampled across 13 African countries revealed no predicted loss-of-function (high-impact) variants within the nAChR gene family. In contrast, synonymous and nonsynonymous mutations were detected, and their allele frequency spectra compared across subunits and countries. All subunits showed synonymous mutations spread in all frequency classes as expected for presumably neutral variants (Fig. 2). In contrast, the distribution and frequencies of nonsynonymous mutations were very heterogenous among subunits. Six subunits (α1, α2, α4, α5, α6, and β1) were highly constrained and showed no nonsynonymous variants. Although the two subunits, α7 and α8 harbored respectively four and one nonsynonymous variants, these mutations were mostly restricted to very low frequency classes (< 0.05). α3 carried three nonsynonymous mutations, including one that reached very high frequencies in several countries. In contrast, the two subunits (β2 and α9) showed signatures of relaxed purifying selection, consistent with the accumulation of numerous missense variants at low, intermediate, and high frequencies. On both subunits, allele frequency spectra of synonymous and nonsynonymous mutations mirrored each other, with variants spanning all frequency classes.

**Figure 2.**
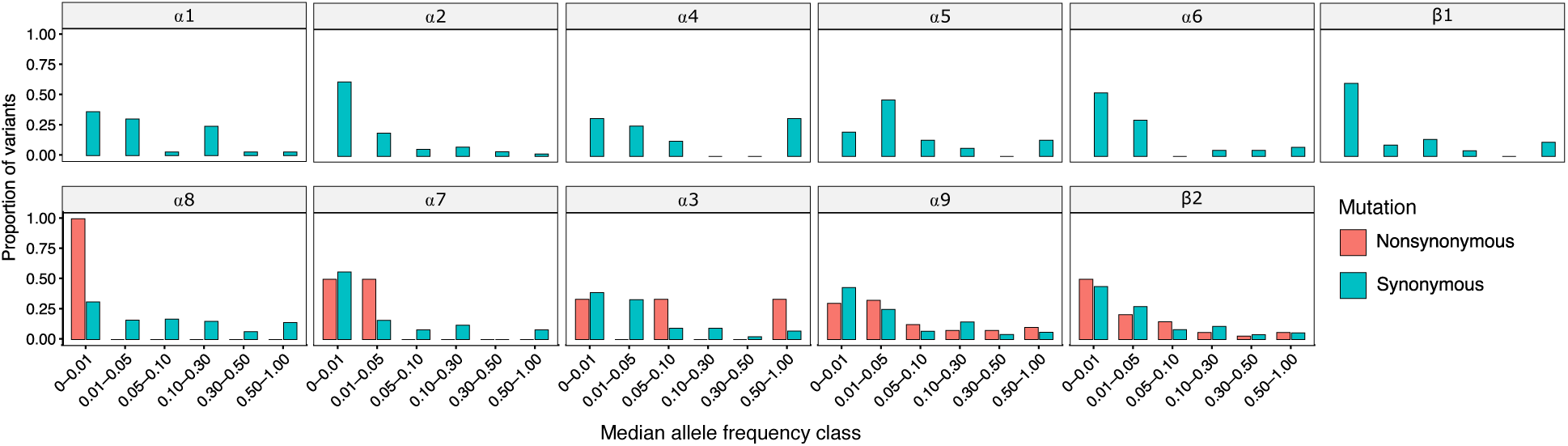
Allele frequency spectra for synonymous and nonsynonymous mutations across *An. funestus* nAChRs. Median allele frequencies were calculated for 656 individuals sampled from 13 African countries. Six subunits are strongly constrained with no nonsynonymous mutations. In the least constrained subunits (α9 and β2), all median allele frequency classes contain both synonymous and nonsynonymous mutations.

Detailed analysis of allele frequencies among countries revealed that nonsynonymous variants can reach intermediate to high frequencies in large geographic areas within the *An. funestus* distribution range (Fig. 3). The single α8 nonsynonymous mutation was a leucine-to-valine change in the C-terminal region and was detected in three countries (Democratic Republic of Congo, Kenya, and Uganda) at a median allele of 0.14 (Inter Quartile Range, IQR 0.12-0.15). In α7, the four nonsynonymous mutations identified including two in the N-terminal region. One of the N-terminal mutations was an isoleucine-to-leucine substitution detected in six countries at a median allele frequency of 0.15 (IQR 0.09-0.35). α3 exhibited a cluster of three nonsynonymous substitutions affecting residues at the intracellular junction between the TM1 and TM2 segments. The most widespread of these was a substitution of lysine to glutamic acid detected in twelve countries, reaching a peak allele frequency of 75% in Central African Republic.

**Figure 3.**
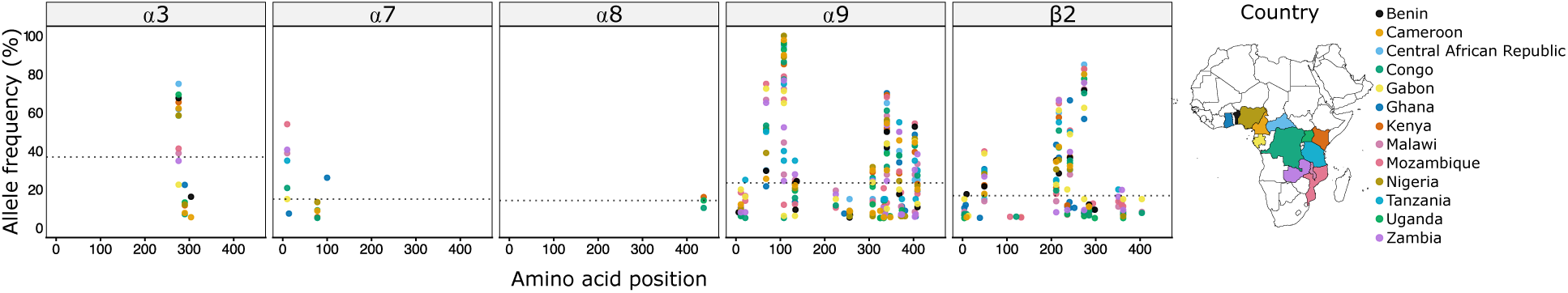
Allele frequency distribution of nonsynonymous mutations across *An. funestus* nAChRs in Africa. Allele frequencies among 13 populations from Africa are shown for the five nAChR subunits carrying nonsynonymous mutations. Each point corresponds to an amino acid substitution at a specific position within a subunit, with colors indicating the country of origin.

Consistent with their evolutionary flexibility, α9 and β2 were the most polymorphic subunits, with numerous nonsynonymous mutations distributed throughout the protein sequence. In both subunits, respectively three and two missense variants had median allele frequencies exceeding 50% throughout the continent. Additionally, intermediate-frequency variants (median allele frequency between 25% and 50%) were detected at four and two residues in α9 and β2, respectively. The most geographically widespread nonsynonymous mutations across the nAChR gene family included two substitutions at that same residue 108 (V108I and V108A) in α9. Both mutations were detected in all 13 sampled countries. V108I was nearly fixed in three countries (Nigeria, Kenya, and Cameroon) and exhibited median mutant-allele frequency of 0.95 (IQR 0.77-0.96) continentwide. V108A had median allele frequency of 0.85 (IQR 0.65-0.93). Other substitutions with median allele frequencies > 50% and present in at least five countries included I340L (0.56, IQR 0.54-0.65) and S68G (0.51, IQR 0.30-0.65) in α9, as well as I274L (0.76, IQR 0.72-0.80) and R217K (0.55, IQR 0.42-0.61) in β2.

### 3.3. Moderate differentiation of nAChR mutations among countries

The single nonsynonymous mutation on α8 exhibited a median *F*_ST_ of 0.10 (IQR 0.05-0.13) among populations throughout the continent (Fig. 4). In α3, the three amino acid substitutions showed low to moderate median *F*_ST_ values ranging from 0.01 (IQR 0.00-0.04) to 0.08 (IQR 0.0009-0.22). However, a glutamate-to-lysine substitution on this subunit was characterized by four pairwise comparisons showing *F*_ST_ values above 0.5, indicating substantial divergence among specific country pairs. This mutation was among the top two most differentiated variants across the nAChR gene family.

**Figure 4.**
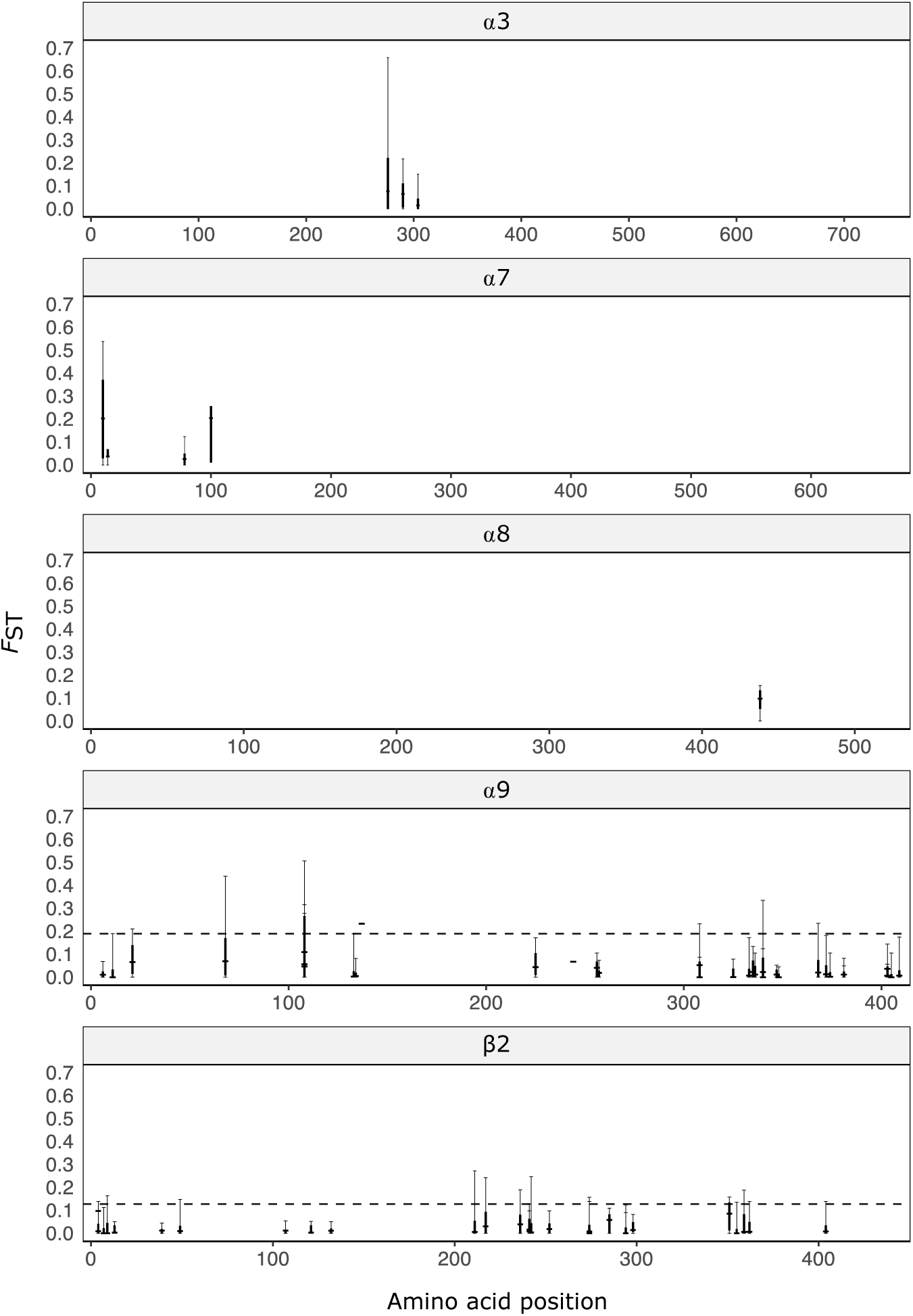
Genetic differentiation (*F*_ST_) of nonsynonymous mutations across nAChR subunits in An. funestus. Pairwise *F*_ST_ values computed for nonsynonymous mutations across five nAChR subunits in 13 *An. funestus* populations are shown. Each point represents the *F*_ST_ value of an amino acid substitution between population pairs, plotted by its position within the protein sequence. Vertical whiskers represent the full observed range of pairwise *F*_ST_ values for each mutation. Thick central bars indicate the interquartile range, and horizontal ticks denote the median *F*_ST_ across country pairs. The dashed horizontal line represents the 95^th^ percentile threshold, highlighting highly differentiated variants.

Among the four nonsynonymous mutations identified in α7, the threonine-to-alanine substitution in N-terminal domain (T100A) displayed the highest differentiation (median *F*_ST_ = 0.20, IQR 0.01-0.26). Four pairwise comparisons revealed *F*_ST_ values above 0.2 consistent with elevated differentiation between countries. Median *F*_ST_ values for α9 mutations were generally below 0.1, reflecting low to moderate geographic structure across the continent. The 95^th^ percentile of the *F*_ST_ distribution for this subunit was 0.09, and only two mutations exhibited multiple pairwise comparisons exceeding this threshold. These mutations included the valine-to-alanine (V108A), which had a median *F*_ST_ of 0.11 (IQR 0.01-0.26) across all pairwise comparisons among countries. The second mutation with multiple pairwise comparisons above the 95^th^ percentile was I137T, which had a maximum *F*_ST_ value of 0.23 detected between Benin and Cameroon.

Similarly, amino acid substitutions on the β2 subunit showed low to moderate overall differentiation, with median *F*_ST_ values per mutation below 0.1 and a 95^th^ percentile threshold of 0.19 (Fig. 4). Among the 28 nonsynonymous mutations detected on this subunit, three had a maximum *F*_ST_ above 0.24: a valine-to-glutamic acid at position 211 (V211E, max *F*_ST_ = 0.27 between Ghana and Zambia), a lysine-to-threonine at position 242 (K242T, max *F*_ST_ = 0.25 between Ghana and Malawi) and an arginine-to-lysine at position 217 (R217K, max *F*_ST_ = 0.24 between Benin and Zambia).

### 3.4. Nonsynonymous mutations in orthosteric ligand-binding and transmembrane domains

We used both sequence-defined domain boundaries and AlphaFold2-based structural models to map amino acid substitutions onto the receptor architecture. The AlphaFold2 models exhibited well-resolved folding and structural accuracy across regions involved in ligand binding and channel formation (pLDDT > 90), supporting their use for structural mapping (Fig. S1, supplemental material). Among *An. funestus* populations, nine amino acid substitutions were detected in ligand-binding domains delimited by sequence motifs in two subunits (β2 and α9) (Fig. 4). In α9, three mutations fell in loops E, C and G. Based on the AlphaFold2-derived pentameric model ((α9)₃(β1)₂), we approximated the orthosteric binding pocket as a predicted envelope within a 6 Å radius surrounding the bound ligand (EPB), corresponding to approximately 5% of the receptor surface (83). As shown in Figures 5 and 6, all three nonsynonymous mutations identified in ligand-binding loops map to the orthosteric domain, with one variant (S68G) located directly within the predicted binding pocket.

**Figure 5.**
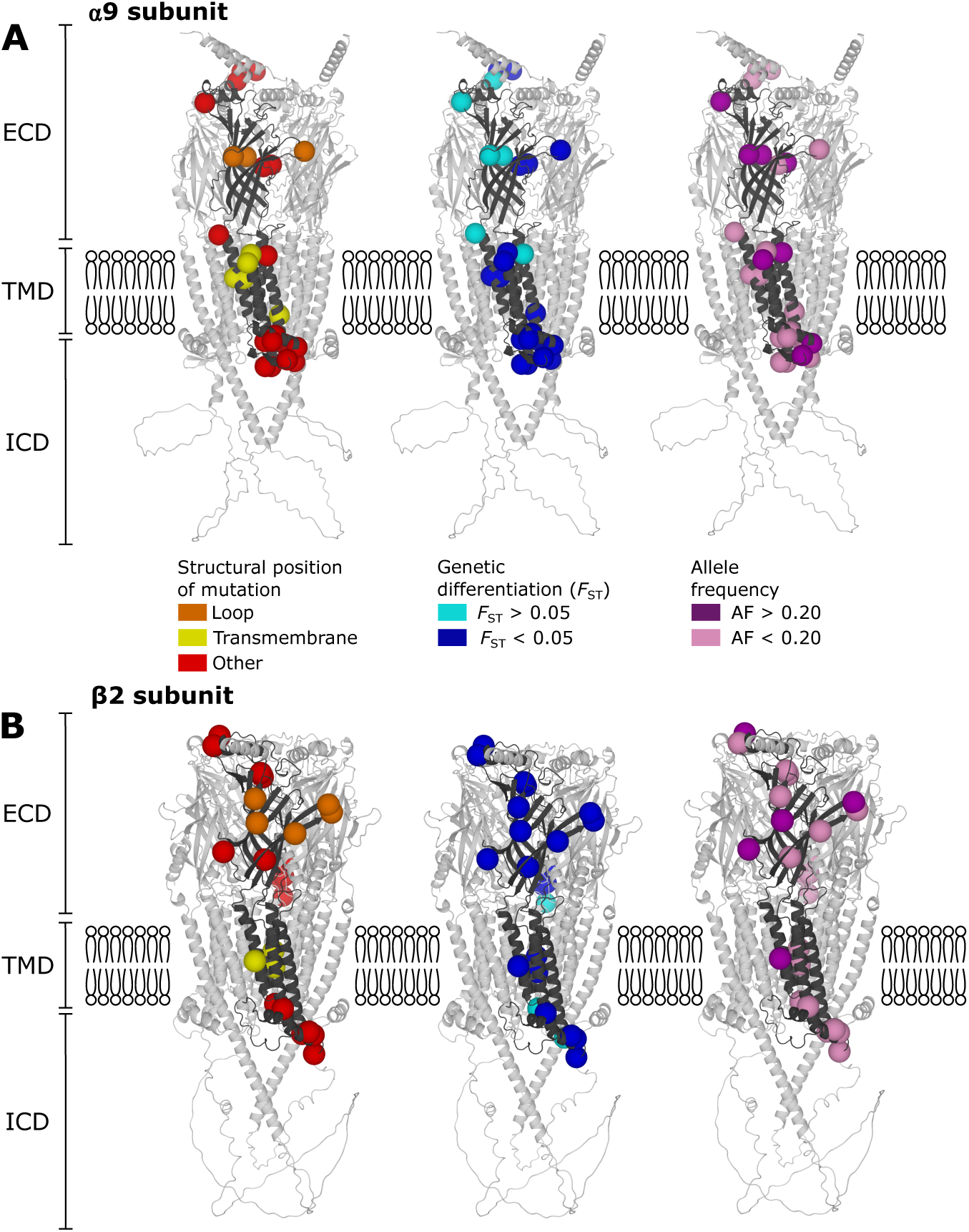
Structural mapping of nonsynonymous mutations in the α9 and β2 nAChR subunits in An. funestus. AlphaFold2-based protein models were assembled into pentameric receptor structures with the α9 (A) and β2 (B) subunits highlighted to visualize the spatial distribution of nonsynonymous mutations. Variants are mapped onto the extracellular (ECD), transmembrane (TMD), and intracellular (ICD) domains. From left to right, residues are colored according to structural location, genetic differentiation, and allele frequency.

**Figure 6.**
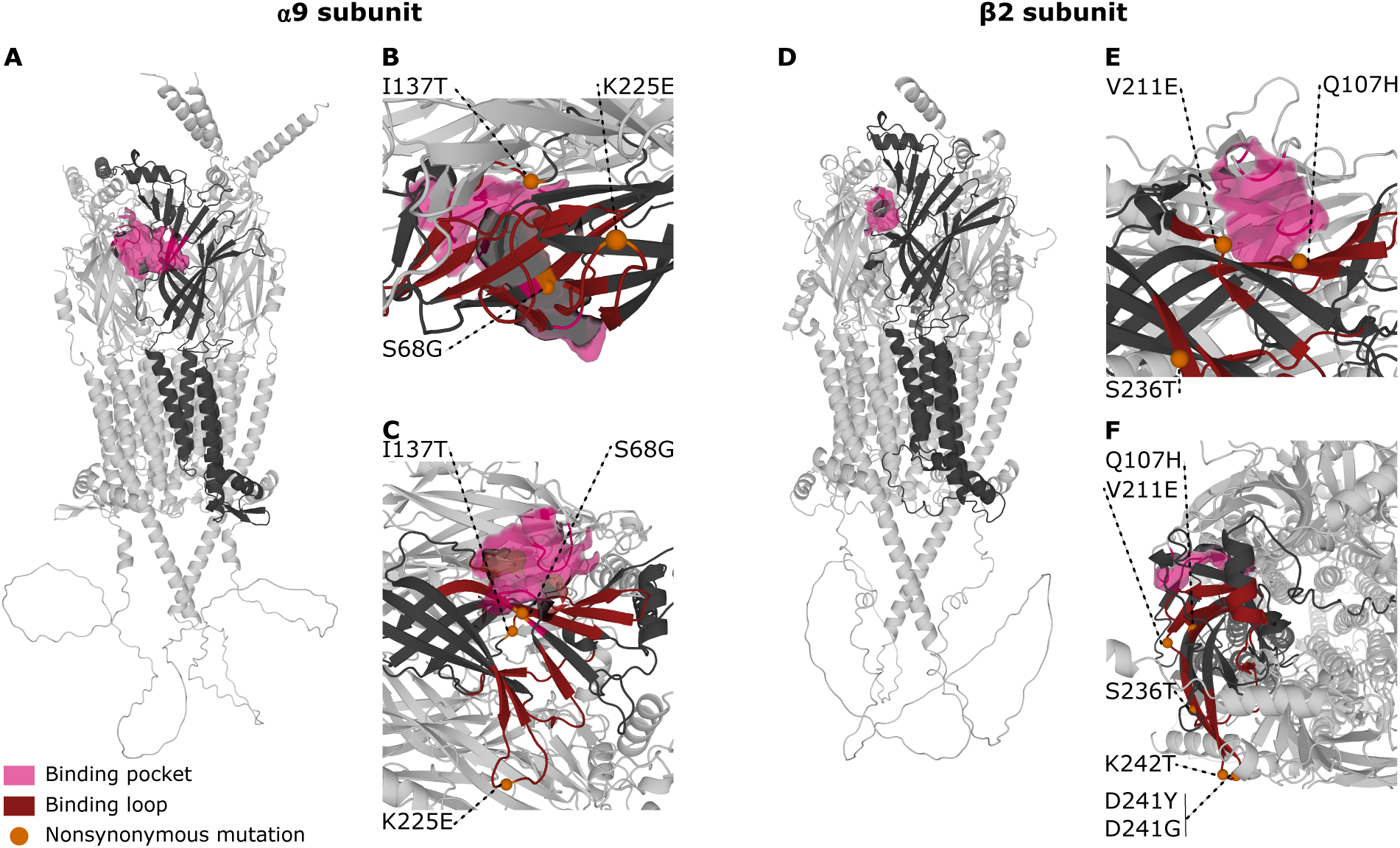
Structural mapping of nonsynonymous mutations within ligand-binding domains of α9 and β2 nAChR subunits in *An. funestus*. Pentameric models were built using AlphaFold2 protein structures, and the nonsynonymous mutations present in orthosteric ligand-binding domains in the α9 (A–C) and β2 (D–F) subunits were highlighted. Top (B and E) and side (C and F) views show the spatial arrangement of the binding pocket and surrounding loops.

The most geographically widespread mutation on the α9 subunit was a serine-to-glycine (S68G) substitution in loop G, detected in 13 countries with a median allele frequency of 0.51 (IQR 0.30-0.65) and a maximum frequency in populations from Mozambique (0.75). An isoleucine-to-threonine (S137T) mutation in loop E was found exclusively in populations from Benin at frequency of 0.24. The lysine-to-glutamic acid change in loop C (K225E) was relatively widespread and occurred in five countries (DRC, Gabon, Malawi, Mozambique, and Zambia) with a continent-wide median allele frequency of 0.15 (IQR 0.12-0.17). The loop-associated α9 mutations exhibited moderate levels of genetic differentiation. The two mutations that appeared in multiple countries showed median pairwise *F*_ST_ of 0.04 (IQR 0.01-0.11) for the K225E in loop C and a median *F*_ST_ of 0.07 (IQR 0.008-0.17) for the S68G in loop G (Fig. 5).

On the β2 subunit, the canonical ligand-binding loops containing nonsynonymous mutations were loops C, D and F. Five mutations were detected within these loops (Fig. 5). The AlphaFold2-derived pentameric model of β2 ((α1)₃(β2)₂) indicated that these mutations mapped to the orthosteric domain, with the V211E and Q107H substitutions located closest to the predicted binding pocket (Fig. 6). Four mutations were detected in loop C, including two that affected the same residues and were restricted to Ghana (D241Y and D241G) as well as a threonine-to-lysine (K242T) substitution at the neighboring residue that was found in all 13 sampled countries. The median allele frequency of this widespread mutation was 0.36 (IQR 0.32-0.49) at the continental scale, with multiple pairwise *F*_ST_ values above 0.20 among countries. In addition, a glutamine-to-histidine substitution in loop D (Q107H) was restricted to Mozambique with a frequency of only 0.06 within the country. The S236T mutation in loop C occurred at low frequency across the continent (median AF 0.10, IQR 0.08-0.12). The final mutation in canonical loops of β2 was a V211E substitution in loop F, which reached a median allele frequency of 0.24 (IQR 0.19-0.30) and was present in all 13 sampled countries. The median pairwise *F*_ST_ value for this mutation was 0.006 (IQR 0-0.06) and reached a maximum of 0.27 between Ghana and Zambia. None of the residues harboring mutations in β2 and α9 binding loops correspond to homologs of canonical residues known to establish electrostatic or hydrogen-bond interactions with neonicotinoids in orthosteric domains.

Across the nAChR gene family, 11 substitutions fell within transmembrane domains (Fig. 5). As for ligand-binding domains, transmembrane mutations were restricted to the two subunits β2 and α9. In α9, substitutions were detected in three transmembrane segments (TM1, TM3 and TM4). In TM1, two amino acid changes (C256S and A257E) affected adjacent residues. The A257E change was restricted to two countries (Benin and Nigeria), whereas C256S was detected in six countries. Both mutations were rare, with median allele frequencies of 0.08 (IQR 0.07-0.08) for A257E and 0.07 (IQR 0.06-0.09) for C256S. Another low-frequency mutation (histidine-to-tyrosine: H325Y) in α9 TM3 was found in Cameroon, Ghana, and Uganda with a median allele frequency of 0.05 (IQR 0.05-0.07) across the three countries. In the TM4 segment, an isoleucine-to-valine substitution at position 403 (I403V) occurred at a median allele frequency of 0.43 (IQR 0.35-0.46) in 13 countries, whereas an isoleucine-to-leucine substitution at the same residue (I403L) was found at low frequency (0.07, IQR 0.06-0.08) in only two countries (Malawi and Zambia). Another low-frequency mutation found in the TM4 segment was a cysteine-to-serine substitution (C405S), detected in four countries at a median allele frequency of 0.07 (IQR 0.07-0.07). The most frequent and most widespread α9 transmembrane mutation, the I403V in TM4, remained moderately differentiated, with pairwise *F*_ST_ values below 0.2 in all comparisons among countries. On the β2 subunit, two substitutions affected the same residue in TM1, resulting in a high-frequency mutation (I274L, median AF: 0.76, IQR 0.72-0.80) widespread in 13 countries and a low-frequency substitution (I274M, median AF: 0.08, IQR 0.07-0.09) detected in seven countries (Fig. 5). In the TM2 segment, which serves as the ion channel, three mutations were detected across the β2 subunit, but they were restricted to one or two countries with median allele frequencies below 0.15. The most frequent mutation in β2 TM2 was an alanine-to-valine substitution at position 294 detected at a frequency of 0.13 among populations from Mozambique.

## 4. Discussion

In this study, we examined genetic variation across nicotinic acetylcholine receptors (nAChRs) in *An. funestus*, a highly efficient malaria vector widely distributed throughout sub-Saharan Africa (69,84,85). With eleven subunits, the nAChR gene family in *An. funestus* lies within the upper range of diversity observed in insects. Comparatively, *Drosophila melanogaster* and the honeybee *Apis mellifera* possesses respectively ten and eleven subunits (36,86).

In *An. funestus*, both α and β subunits had very high sequence similarity with their orthologs from the distantly related species *An. gambiae* (8,35,72,73). The β2 subunit was the least well conserved with only 73% similarity between species. Multiple phylogenetic studies have identified β2 and α9 as divergent subunits with a greater evolutionary rate than the rest of the nAChR family in insects (36,86,87). All Subunit-encoding genes except three (α5, α9, and β2) are located on homologous chromosomal arms in *An. funestus* and *An. gambiae*. The localization of othologous nAChR subunits on different chromosomes is not uncommon and has been reported for example between *An. gambiae* and *D. melanogaster* due to ancestral chromosomal rearrangements (35,36).

This study revealed evolutionary constraint on most nAChR genes in *An. funestus* with core subunits carrying no nonsynonymous mutations while harboring synonymous variants at intermediate or high frequencies. Such signatures are consistent with purifying selection which purges potentially deleterious nonsynonymous variants and have been observed in nAChRs of two other major malaria mosquitoes: *An. gambiae* and *An. coluzzii*. In both species, only three nonsynonymous variants with frequencies exceeding 5% were detected among 11 nAChR subunits throughout Africa (72). nAChR genes are both essential in neural transmission and pleiotropic with diverse roles in multiple physiological pathways, two criteria known to reduce the evolutionary rate of proteins (88–90). *Anopheles funestus* is unique among malaria vector species because despite the strong evolutionary constraint, some nAChR genes are accumulating missense mutations and evolving rapidly. It has long been known that α9 and β2 are divergent subunits in insects, but the mechanisms maintaining such divergence have remained obscured (91). Here we showed that on both subunits, allele frequency spectra of synonymous and nonsynonymous mutations are comparable indicating signatures of relaxed purifying selection. The specific physiological roles of α9 and β2 in insects remain unclear, but their evolutionary flexibility suggests that they may be less essential and less pleiotropic than core subunits of the nAChR family.

Since nAChR subunits are not isolated proteins and act as units within multimeric assemblies, the elevated rate of evolution in α9 and β2 may serve to maintain a level of diversification of receptors necessary to respond to diverse environmental pressures. Relaxed constrained on subunits, particularly α9 and β2, may provide repositories of standing variation, modulating orthosteric and allosteric interactions, post-transcriptional regulation, and channel gating kinetics within the family (41,42). The fact that some amino acid substitutions display moderate to elevated genetic differentiation among countries also suggests that some of these variants may contribute to geographically structured responses to local selection pressures. Furthermore, the hypothesis of relaxed constraint predicts that nonsynonymous mutations with weak to moderate deleterious effects may persist and occasionally reach intermediate or high frequencies because they are not efficiently removed by purifying selection. However, episodes of positive selection acting on individual alleles that alter the structure and function of pentameric assemblies and confer adaptive advantages under specific geographic or physiological contexts cannot be excluded.

Amino acid substitutions that modulate nAChR affinity to neonicotinoids or spinosyns have been widely documented in agricultural insect pests and represent a canonical mechanism of resistance evolution (55–59,92). However, studies evaluating associations between candidate target-site mutations and insecticide resistance rarely consider the broader mutational landscape of the nAChR gene family. Consequently, it remains unclear whether rapid evolution of specific subunits contributes to insecticide sensitivity beyond the established pharmacological effects of individual point mutations.

Most resistance-associated substitutions detected so far within nAChRs in insects occur in ligand-binding loops of the β1 subunit (59). This subunit is among the most constrained nAChR genes in *An. funestus*, with no amino acid variants across the species’ distribution range in Africa. Knockout of β1 in *D. melanogaster* has be associated with extremely low survival rates indicating an essential gene that likely remains under strong evolutionary constraint unless subjected to intense selection pressures typical of agricultural settings (38).

Other variants associated with insecticide resistance have been reported across α1, α2, α3, and α6 in diverse insect pests (55,93–95). In *An. funestus*, no common nonsynonymous variants were detected in α1, α2, or α6, whereas α3 harbored three amino acid substitutions. Ligand affinity tests have shown that receptor assemblies containing α3 can bind neonicotinoids in both *D. melanogaster* and *An. gambiae* (38,39,61). However, the substitutions identified in this study were located outside canonical orthosteric binding loops suggesting limited potential for directly altering neonicotinoid affinity in *An. funestus*. In addition, none of the α3 variants corresponded to the Y151S substitution which has been shown to confer resistance to imidacloprid in *Nilaparvata lugens* (55).

Five nonsynonymous mutations including one geographically widespread variant were detected outside orthosteric domains of α7 and α8 in *An. funestus*. Neither subunit is commonly implicated in neonicotinoid resistance in agricultural pests, and knockdown tests conducted in *D. melanogaster* suggest that both subunits have lower neonicotinoid affinity compared to β1 (38,39).

The two divergent subunits across insects (α9 and β2), which are also the most polymorphic in *An. funestus* as revealed by our population genomics study, exhibit relatively low direct affinity for several neonicotinoid insecticides compared to β1, α1, or α3 for instance (38). Nevertheless, both α9 and β2 carry amino acid substitutions at intermediate to high frequency in orthosteric binding domains which contain basic residues critical for insecticide affinity. Within the α9 subunit, mutations affecting the orthosteric region were observed in loops C, E, and G. Loop C of α subunits contains residues capable of forming electrostatic interactions with neonicotinoids, as demonstrated through mutagenesis and electrophysiological studies (48–50). In *D. melanogaster* for instance, a conserved proline residue in loop C contributes to neonicotinoid binding (96). The only loop C nonsynonymous mutation detected in *An. funestus* in this study was a lysine-to-glutamic acid change at position 225, while the proline described in *D. melanogaster* remained monomorphic across continental populations. Mutagenesis experiments in *Nilaparvata lugens* have shown that some substitutions in loop E (S131Y/R and D133N) can alter neonicotinoid sensitivity (46). In contrast, we detected a single mutation in loop E of *An. funestus* restricted to one country: an isoleucine-to-threonine transition whose potential role in binding remains to be determined. In the β2 subunit, orthosteric substitutions were identified in loops C, D, and F. Loop D is of particular functional importance in β subunits because it contains a conserved arginine residue critical for neonicotinoid binding (45,96). Aside from a low-frequency glycine-to-histidine substitution (mean MAF < 6% continent-wide), no nonsynonymous variation was detected in loop D in *An. funestus*.

Transmembrane domains in cys-loop receptors are essential to channel gating as well as ion conductance and selectivity. These domains are therefore expected to be relatively constrained, yet nonsynonymous mutations including high frequency variants were detected across all four transmembrane domains (M1–M4) of α9 and β2. Because these segments play distinct structural and functional roles, their mutations may have variable effects on receptor properties (27,41–43). The M1, M3, and M4 helices contribute primarily to receptor stability and interactions with the lipid environment, whereas the M2 segment forms the pore-lining helix that controls channel gating and cation selectivity (42,97). As the ion channel, M2 can be considered more essential to the receptor functioning and subjected to stronger evolutionary constraint. Three low- to moderate-frequency substitutions were detected within the M2 segment of β2 and were geographically restricted to one or two countries. Although their functional implications remain to investigated, amino acid changes that modify hydrophobic interactions, such as the alanine-to-valine substitution (A294V), may have greater potential to alter ion conductance or selectivity (42).

The observed replacement of *An. gambiae* by *An. funestus* in some areas following the implementation of clothianidin-based indoor residual spraying has raised the possibility that *An. funestus* has greater intrinsic capacity to tolerate or develop resistance to this active ingredient (4,5). While this study highlights amino acid variation that could alter *An. funestus* sensitivity to neonicotinoids, insect susceptibility to insecticides reflects multiple physiological and behavioral mechanisms that were not addressed here (14,25,98,99). In particular, metabolic detoxification, behavioral adjustments, and differential expression of nAChR subtypes are associated with reduced neonicotinoid susceptibility, but the extent to which these processes contribute to adaptive responses in *An. funestus* remains to be determined (61,62,99–101).

## Conclusions

The rapid emergence of clothianidin resistance in *An. funestus* underscores the importance of proactive resistance management to extend the operational lifespan of newly deployed vector control products (4,5). This study provides a family-wide assessment of molecular diversity across nAChRs in *An. funestus* and establishes a baseline for monitoring the emergence and spread of resistance-associated mutations as insecticide pressure intensifies. Genomic surveillance, together with functional validation of nAChR mutations, will be essential for the early resistance detection and for informing sustainable deployment strategies for neonicotinoid-based interventions in vector control.

## Ethical approval

Not applicable.

## CRediT authorship contribution statement

**Desiree Rios**: Formal analysis, Investigation, Methodology, Writing–review and editing. **Caroline Fouet**: Conceptualization, Formal analysis, Investigation, Methodology, Writing–review and editing. **Colince Kamdem**: Conceptualization, Formal analysis, Investigation, Methodology, Supervision, Writing–original draft, Writing–review and editing.

## Data availability

Variant call format (VCF) files from the *Anopheles funestus* genomic surveillance project analyzed in this study are available from the MalariaGEN repository (https://www.malariagen.net/project/anopheles-funestus-genomic-surveillance-project/). Accession numbers of the 656 sequenced samples are provided in (69).

## Figure legends

**Figure S1.**
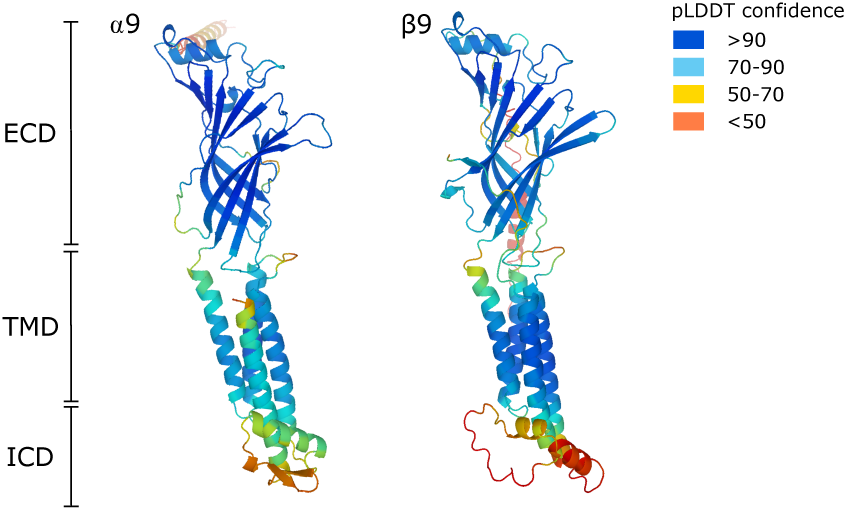
Confidence scores of AlphaFold2 models for α9 and β2 nAChR subunits of *Anopheles funestus*. AlphaFold2 structural models of the α9 and β2 subunits are colored according to predicted Local Distance Difference Test (pLDDT) scores which provide an estimate of how accurately the model predicts the local environment of each amino acid. The extracellular domain (ECD) and transmembrane domain (TMD) exhibit high confidence (pLDDT > 90), indicating well-resolved folding and structural accuracy in regions involved in ligand binding and channel formation. Confidence scores are lower in flexible linker regions, while pLDDT score < 50 are restricted to a small portion of the intracellular domain (ICD).

